# Neural signatures of reading-related orthographic processing in braille

**DOI:** 10.1101/2022.11.09.515790

**Authors:** Yun-Fei Liu, Brenda Rapp, Marina Bedny

## Abstract

Blind readers use a tactile reading systems consisting of raised dot arrays: braille/⠃⠗⠇. How does the human brain implement reading by touch? The current study looked for signatures of reading-specific orthographic processes in braille, separate from low-level somatosensory responses and semantic retrieval. Of specific interest were responses in posterior parietal cortices (PPC), because of their role in high-level tactile perception. Congenitally blind, proficient braille readers read real words and pseudowords by touch while undergoing fMRI. We leveraged the system of contractions in English braille, where one or more braille cells can represent combinations of English print letters (e.g., “ing” ⠬, “one” ⠐⠕), making it possible to separate physical and uncontracted letter-length. All words in the study consisted of 4 braille cells, but their corresponding Roman spellings varied from 4 to 7 letters (e.g., “con-c-er-t” ⠒⠉⠻⠞. contracted: 4 cells; uncontracted: 7 letters). We found that the bilateral supramarginal gyrus (SMG) in the PPC increased its activity as the uncontracted word length increased. By contrast, in the hand region of primary somatosensory cortex (S1), activity increased as a function of a low-level somatosensory feature: dot-number per word. The PPC also showed greater response to pseudowords than real words and distinguished between real and pseudowords in multi-voxel-pattern analysis. Parieto-occipital, early visual and ventral occipito-temporal, as well as prefrontal cortices also showed sensitivity to the real-vs-pseudoword distinction. We conclude that PPC is involved in sublexical orthographic processing for braille, possibly due to braille’s tactile modality.

**Significance statement:** Blind readers use tactile reading systems of raised dot arrays: braille. To identify signatures of orthographic processing for reading by touch, and dissociate it from tactile and linguistic process, we leveraged the system of contractions in English braille, where one or more braille characters represents combinations of English print letters. Blind proficient braille readers read real words and pseudowords during fMRI scans. While all words consisted of 4 braille characters, the uncontracted spelling ranged from 4-7 letters. Activity in bilateral-posterior-parietal cortices, just posterior to primary-somatosensory cortex, increased with uncontracted word length, independent of tactile complexity (number of raised dots per word). By contrast, primary-somatosensory activity increased with tactile complexity. The posterior-parietal cortices contribute to tactile reading.

## Introduction

The invention of writing approximately 5,000 years ago transformed human communication and enabled both technological and cultural innovation. How does the human brain enable reading, a cultural invention for which it could not have evolved dedicated mechanisms? Cognitively, reading engages a collection of orthographic processes, including letters/symbol recognition, retrieving stored representations of the spellings of familiar words from the orthographic (long-term memory, LTM) lexicon (supporting the reading of irregular words e.g., colonel) and grapheme-to-phoneme conversion (i.e., connecting sublexical orthographic units, such as letters and digraphs [Bà /b/, etc.]), to speech sounds (supporting the reading of novel words, e.g., glorfomistic) (Fischer-Baum & Rapp, 2014; Forster & Chambers, 1973; Marshall & Newcombe, 1973). Remarkably, despite its recent cultural invention, these reading related processes engage similar neural circuits across a variety of languages and writing systems, including alphabetic (e.g., English, French), logographic (e.g., Chinese), and syllabic (e.g., Japanese kana) visual scripts (Bai, Shi, Jiang, He, & Weng, 2011; Dehaene et al., 2010; Liu et al., 2008; Nakamura, Dehaene, Jobert, Le Bihan, & Kouider, 2005; Nakamura et al., 2012; Rueckl et al., 2015).

With the acquisition of literacy, a region in the mid-fusiform gyrus of the ventral occipito-temporal cortex (vOTC) becomes specialized for processing written letters, letter combinations and words in reading as well as spelling (Dehaene & Cohen, 2007; Dehaene, Le Clec’H, Poline, Le Bihan, & Cohen, 2002; McCandliss, Cohen, & Dehaene, 2003; Purcell, Turkeltaub, Eden, & Rapp, 2011). Because of its importance in reading, this part of the vOTC has been often referred to as the “visual word form area (VWFA)” (Dehaene et al., 2002). Other cortical areas that are important for reading include left-lateralized regions in inferior frontal and posterior parietal cortices (PPC), with the latter implicated in orthographic working memory maintenance during effortful reading (Cohen, Dehaene, Vinckier, Jobert, & Montavont, 2008; Deschamps, Baum, & Gracco, 2014; Graves, Desai, Humphries, Seidenberg, & Binder, 2010; Price, Wise, & Frackowiak, 1996).

Most of the world’s reading systems make use of visual symbols and most of what we know about the neural basis of reading concerns visual reading. By contrast, braille is a distinctive tactile orthographic system used primarily by blind and visually-impaired readers whose neural bases are still poorly understood. Braille is read by passing the fingers across a raised dot array. Each braille symbol, called a “cell”, is constructed out of six possible dot positions arranged in a 3(horizontal)-by-2(vertical) array. Apart from being tactile, braille differs from print in that it consists of disconnected dots and therefore lacks lines or line junctions, which have been noted as a universal property of visual scripts (C. H. Chang et al., 2015; Dehaene, Cohen, Sigman, & Vinckier, 2005). Although raised print letters with lines were in use in 18^th^ and 19^th^ century, braille proved to be a more efficient tactile reading code and has been universally adopted by blind communities all over the world after its invention by a blind 15-year-old student, Louis Braille in 1829.

The form of Braille most commonly used by English-speaking proficient blind readers is called Contracted Unified English braille (UEB) (Simpson, 2013). Contracted UEB braille contains 26 cells designated to represent each Roman letter of the English alphabet (e.g., “⠁” represents “a”, “⠃” represents “b”, and “⠵” stands for “z”). However, some Roman letter combinations are represented with strings of one or more braille cells called “contractions”. The number of braille cells in a contraction is always less than the number of Roman letters it represents. For example, “ing” is represented with a single cell “⠬”, “er” is represented with another single cell “⠻”, and the string “-one-” in the word “honey” (⠓⠐⠕⠽) is represented by a two-cell contraction “⠐⠕”. These are examples of contractions with their own braille cells or cell combinations, while other contractions may be represented by multi-purpose single cells that serve to represent either a single letter or a word, depending on the context. For example, ⠓ stands for the letter “h” when it is part of a word like “happy” (⠓⠁⠏⠏⠽), but when presented alone, it stands for the word “have”. Since contractions stand for frequent letter combinations and words, they are an important part of naturalistic English braille reading.

Blind English-speakers learn the spellings of words in the uncontracted (i.e., Roman letter) English alphabet as well as the contracted forms of words and the contraction rules (Millar, 1997). Proficient English-speaking readers most frequently use contracted braille to read, and for writing in braille directly on a Perkins brailler or a smartphone. However, uncontracted spellings are typically used when writing on a computer keyboard or when spelling out loud. Proficient English braille readers thus know two orthographic codes for spelling in braille: contracted and uncontracted.

The cognitive and neural interactions of contracted and uncontracted braille codes during reading are not known. Proficient braille readers are faster to read contracted versions of frequent words than their uncontracted, letter-by-letter spelled out counterparts (e.g., “s-u-g-ar” contracted ⠎⠥⠛⠜; uncontracted ⠎⠥⠛⠁⠗, “s-u-g-a-r”) and the task of matching uncontracted, and therefore uncommon, versions of frequent words to their contracted counterparts is effortful (Millar, 1997). However, when contractions interrupt the sub-lexical structure of a word, they slow down braille reading. That is, contractions that span across boundaries of sub-lexical units such as morphemes (e.g., ⠗⠢⠑⠺, r-**en**-e-w) are read more slowly than contractions that do not interrupt morphemes (e.g., ⠢⠞⠗⠽, **en**-t-r-y) (Fischer-Baum & Englebretson, 2016). This finding is analogous to what has been reported for print reading, where it takes longer to read a word if it has been artificially segmented in a way that violates the boundaries of the sub-lexical linguistic units, than if the segmenting does not violate those units (e.g., for French readers, segmenting the word “champignon” in to “ch-am-p-i-gn-on” is easier to read than “c-ha-mp-igno-n”) (Bouhali, Bézagu, Dehaene, & Cohen, 2019; Prinzmetal, Treiman, & Rho, 1986; Rapp, 1992).

The distinctive nature of braille as a writing system makes understanding its neural basis of particular interest, yet much remains unknown about how braille is neurally implemented. Previous studies have identified a wide network of areas that are active while reading braille. The anatomical location of the visual word form area (VWFA), which is important for visual print reading, is also active during braille reading (Büchel, Price, & Friston, 1998; Burton, Sinclair, & Agato, 2012; Rączy et al., 2019; Reich, Szwed, Cohen, & Amedi, 2011; Sadato et al., 1998). However, the VWFA location has been shown more responsive to speech and high-level linguistic information (e.g., grammatical complexity of spoken sentences) in blind than in sighted individuals, although some responses to spoken language in this region are observed in the sighted (Dzięgiel-Fivet et al., 2021; Kim, Kanjlia, Merabet, & Bedny, 2017; Tian, Saccone, Kim, Kanjlia, & Bedny, 2022). These findings suggest that the VWFA of blind braille readers may not be the main locus for orthographic processing. In addition, braille reading also engages many other occipital regions besides the vOTC, including primary visual cortex (V1) and dorsal occipital cortices (Büchel et al., 1998; Burton, Diamond, & McDermott, 2003; Burton et al., 2012; Burton, Snyder, Diamond, & Raichle, 2002; Sadato et al., 1998).

As a tactile activity, reading braille engages the somatosensory cortices. It also engages the posterior parietal cortices (PPC), which is implicated in high-level tactile perception, as well as parieto-occipital and dorsal occipital cortices posterior to PPC (Büchel et al., 1998; Burton et al., 2003; Burton et al., 2012; Burton et al., 2002; Sadato et al., 1998). While the PPC is sometimes reported in studies of visual reading (Ischebeck et al., 2004; Jobard, Crivello, & Tzourio-Mazoyer, 2003; Vogel et al., 2013; Vogel, Miezin, Petersen, & Schlaggar, 2012), responses to braille are more extensive and more selective for written words (Tian et al., 2022). Moreover, the lateralization of responses to braille words in PPC of blind readers is also suggestive of a reading-related orthographic role, since it is influenced both by reading hand (right or left) and spoken language lateralization (Tian et al., 2022).

Which, if any, of the cortical areas responsive to braille specifically support reading-specific orthographic processes, as opposed to tactile discrimination and semantic retrieval, remains unclear. Most prior studies have compared braille reading to low level control conditions that differ from braille in somatosensory and linguistic properties, and these studies have relied exclusively on univariate analyses (Burton et al., 2012; Kim et al., 2017; Rączy et al., 2019). Despite contractions being a major part of naturalistic English braille reading, to our knowledge no prior study has specifically examined neural responses to braille contractions. The objective of the current experiment was to look for neural signatures of reading-specific form-based orthographic processing in braille and to separate these from low-level somatosensory and high-level language (e.g., semantic) processing using univariate and multivariate analytic approaches.

Our first goal was to distinguish orthographic from lower-level tactile processes by leveraging braille contractions present in English braille. Recall that English braille uses contractions, where one or more braille cells can represent whole words or strings of letters. For example, the letter string “con” is represented by the braille cell “⠒” (when “con” is the initial syllable of a multi-syllable word). It is therefore possible to study neural effects of the length of corresponding uncontracted forms of words while holding the number of cells in the word constant. For example, the words “c-o-r-n” (⠉⠕⠗⠝) and “con-c-er-t” (⠒⠉⠻⠞) are identical in numbers of braille cells, (i.e., both are 4-cells long) but their corresponding uncontracted forms differ in length (i.e., “concert” is 7 while “corn” is 4 letters long because the former has two contractions). We reasoned that cortical areas involved in orthographic processing would show sensitivity to the length of the underlying uncontracted forms of words, even when the number of cells in the contracted form is held constant.

On the other hand, to look for low-level tactile effects, we quantified the amount of somatosensory stimulation in a given word by counting the number of raised dots. A letter like “a” (⠁) consists of one dot, while “y” (⠽) has five. This physical property of letters is orthogonal to orthography since each letter stands for a unique grapheme, regardless of its dot number. We predicted that words with a greater number of dots would be associated with greater activity in the hand region of early somatosensory cortex, while uncontracted word-length should affect neural activity in orthographic regions of the braille-reading network, outside the primary somatosensory cortex (S1), possibly the PPC.

We also investigated the effect of word frequency on neural activity in braille reading. Less frequent visual and braille words are read more slowly, we tested for a neural signature of frequency in braille (Brysbaert, Mandera, & Keuleers, 2017; Millar, 1997; Monsell, Doyle, & Haggard, 1989). In sighted readers, frequency affects neural activity in different cortical networks than those affected by word length (in Roman letters). We hypothesized that the same would be true for braille (Dehaene & Cohen, 2011; Kronbichler et al., 2004; Kuo et al., 2003; Lin, Yu, Zhao, & Zhang, 2016; Rapp & Dufor, 2011; Rapp, Purcell, Hillis, Capasso, & Miceli, 2016; Schuster, Hawelka, Hutzler, Kronbichler, & Richlan, 2016; Woolnough et al., 2021).

A second goal of the current study was to distinguish form-based orthographic/reading specific processing from higher-level language (i.e., semantic) processing that are also engaged during reading of real words by comparing words to pseudowords. In the case of visual print reading, previous studies find that particularly during slow-presentation tasks, form-based (orthographic and phonological) cortical networks respond more to pseudowords than to real words (Bouhali et al., 2019; Dietz, Jones, Gareau, Zeffiro, & Eden, 2005; Hagoort et al., 1999; Liu et al., 2008; Mano et al., 2012). By contrast, even in attentionally demanding tasks, cortical areas involved in semantic processing respond more to meaningful words (Hagoort et al., 1999; Price et al., 1996; Protopapas et al., 2016; Taylor, Rastle, & Davis, 2013). We predicted that cortical areas sensitive to orthographic form in braille, and in particular, orthographic processing of sub-lexical units, would respond more to pseudowords than real words during braille reading, whereas cortical areas involved in semantic processing would respond more to real words than pseudowords (Kronbichler et al., 2004; Mechelli, Gorno-Tempini, & Price, 2003).

Additionally, we used multivariate pattern analysis (MVPA) to identify cortical networks whose patterns of activity differentiate real and pseudo-words, even once univariate signal differences are controlled. To our knowledge no prior studies have used an MVPA classification approach to examine orthographic/lexical information in patterns of activity during braille reading. Instead of asking whether each single voxel or vertex in the brain is more active in some condition than others, MVPA allows us to ask whether patterns of activity within a cortical area are sensitive to the lexical status of braille stimuli (Haxby, Connolly, & Guntupalli, 2014; Norman, Polyn, Detre, & Haxby, 2006). Even if a cortical area is involved in processing both real and pseudo-words to a similar degree, as measured by univariate responses, multivariate patterns may pick up on distinctive processes and/or representations engaged by real and pseudowords. Areas showing the following three effects: sensitivity to uncontracted word length, univariate sensitivity to pseudowords compared to words, and multivariate differences between pseudo and real words, are particularly good candidates for important neural circuits in form-based orthographic processing of braille.

Finally, we sought, for the first time, to detect unique neural signatures of individual braille words. Since braille reading is a tactile, temporally extended process, this may pose special challenges for fMRI-based analysis. Our approach and exploratory findings may advise future studies on braille reading.

## Method

### Participants

Twelve congenitally blind native English speakers (6 females, M=39.5 years of age, SD=11.3) participated in the study. All of the participants were fluent braille readers who began learning braille in early childhood (M=4.3 years of age, SD=1.5; see Table 1 for the cause of blindness). Since proficient blind braille readers are a small population, we compensated for the relatively small number of participants by collecting a large amount of data per subject (i.e., one hour and forty-five minutes of functional braille reading data per person). In previous studies, we have found that with this approach we can reliably measure strong and reliable braille responses with as few as 10 participants (Kim et al., 2017).

**Table 1.** Participant information

| **Blindness etiology** | **N** |
| --- | --- |
| Leber congenital amaurosis | 5 |
| Retinopathy of prematurity | 4 |
| Born without optic nerve | 1 |
| Optic nerve detached | 1 |
| Unknown retinal defect | 1 |

All of the participants had at most minimal light perception since birth and blindness due to pathology anterior to the optic chiasm. Participants were screened for cognitive and neurological disabilities through self-report. The structural images of the participants were inspected by radiologists in Johns Hopkins Hospital and no gross neurological pathologies were detected. Participants gave informed consent and were compensated $30 per hour. All procedures were approved by the Johns Hopkins Medicine Institutional Review Board.

### Stimuli and Task

Participants read real words (n=60) and pseudowords (n=30) presented on an MRI compatible braille and tactile graphic display consisting of an array of 32 by 35 pins (spaced 2.4 mm apart) (Piezoelectric Tactile Stimulus Device developed by KGS Corporation, Japan, see Bauer et al. (2015)). Each word/pseudoword was comprised of exactly 4 braille cells and was presented in contracted braille.

The English braille code has gone through many revisions. All stimuli in the current experiment were written in UEB, the most current braille system (Simpson, 2013). Real words consisted of inanimate words (e.g., corn, n=30) and animate words (e.g., dancer, n=30) selected from the top 20,000 most frequent words in the Corpus of Contemporary American English (COCA; Davies (2008) https://www.english-corpora.org/coca/) (see Table 2 and Table 3 for stimuli). All pseudowords were created by retaining the first two cells of an inanimate real word and replacing the last two cells with other braille cells, including cells that stand for contractions. For example, the real word ⠒⠧⠑⠽ (con-v-e-y) was modified to create the pseudoword ⠒⠧⠥⠅(con-v-u-k). Prior to the fMRI experiment we asked a braille reader to read aloud the pseudowords and confirm that all of them were pronounceable (see Table 4 for pseudoword stimuli).

**Table 2.**
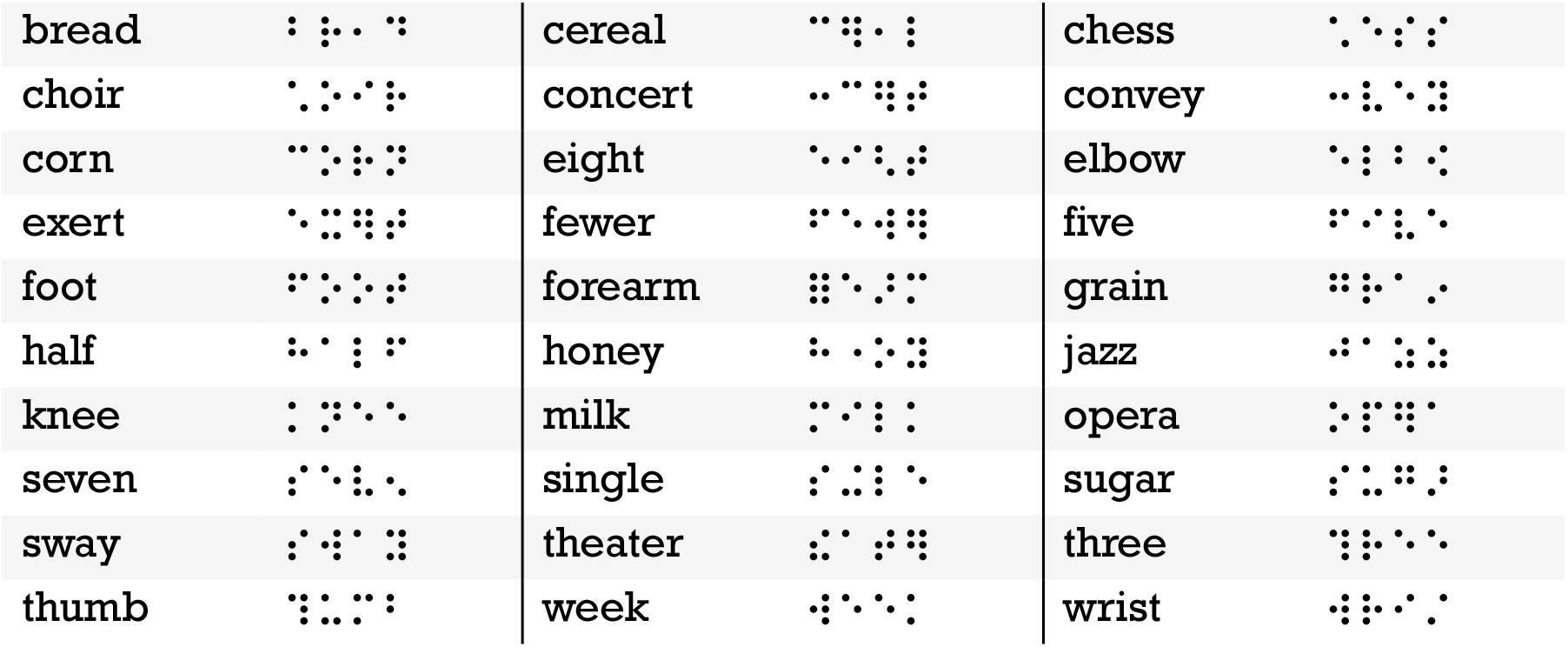
List of inanimate real words and their Braille transcriptions

**Table 3.**
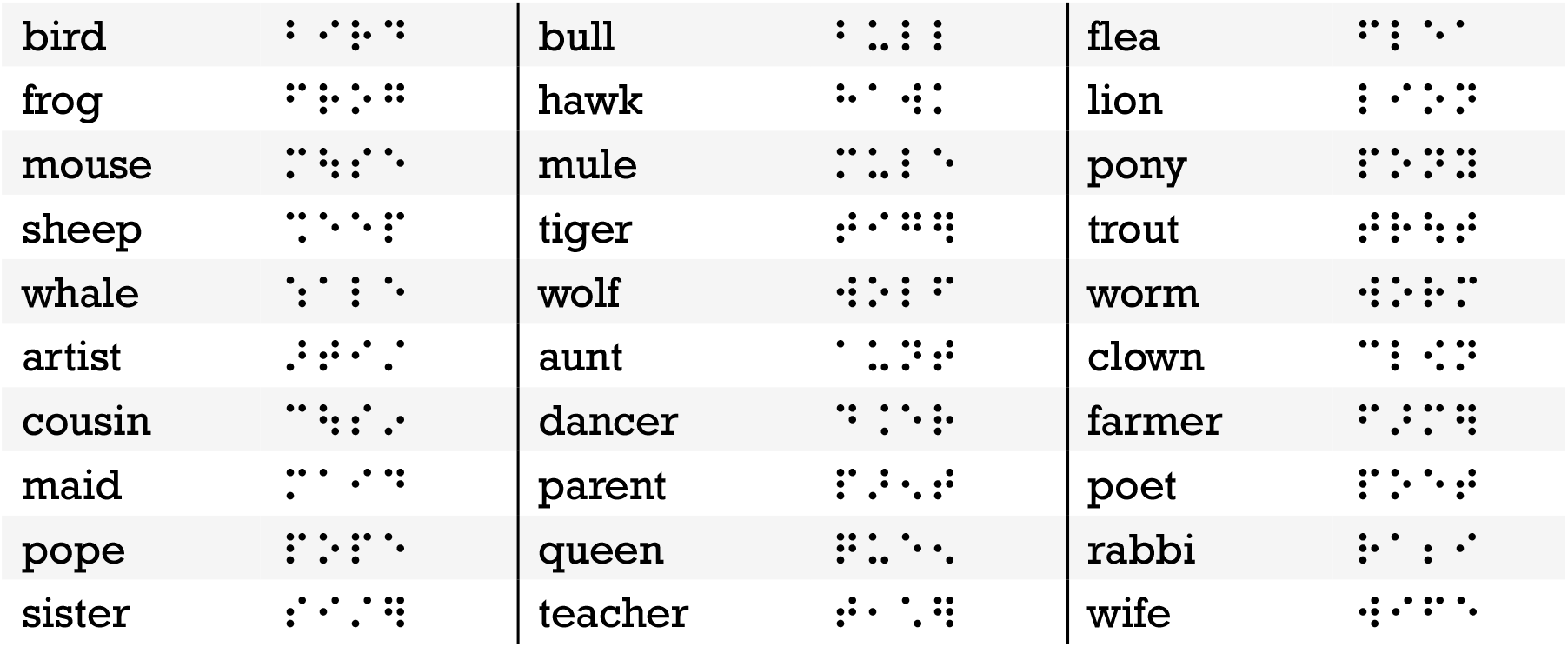
List of animate real words and their Braille transcriptions

**Table 4.**
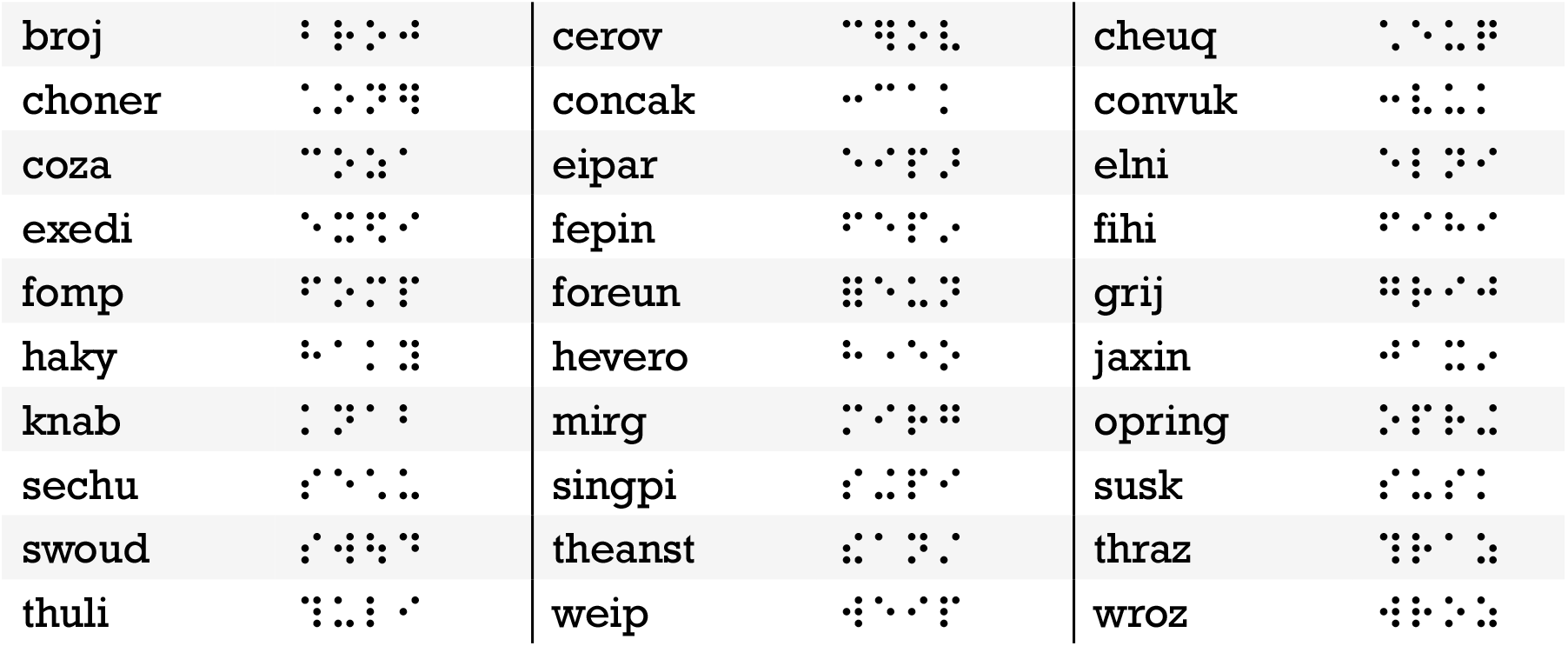
List of pseudowords and their Braille transcriptions

Each trial lasted for 6 seconds, with the braille word displayed for the first 3 seconds and the braille display went blank for the latter 3 seconds. Participants pressed one of three buttons to indicate whether the presented word was an inanimate real word, an animate real word, or a pseudoword. They were instructed to make a response as soon as they knew the answer. We chose this dual task procedure to ensure that participants were attending both to the orthography and the meaning of the words.

The experiment consisted of 15 runs (participants were free to take breaks between runs). In each run, all 30 inanimate real words were presented twice in different orders, and 4 distinct animate words and 4 distinct pseudowords were interspersed throughout the run in pseudo-random intervals. On average, there was 1 animate real word or pseudoword every 7 target words and animate real words and pseudowords were never presented consecutively. Throughout the whole experiment, each inanimate real word was presented 30 times, and each animate real word and each pseudoword was presented twice.

During the fMRI experiment, participants were instructed to read with the index finger of their preferred reading hand, and to press the response buttons using the other hand. Six of the twelve participants read with their right hands.

### fMRI acquisition and preprocessing

All functional and structural MRI data were acquired at the F.M. Kirby Research Center of Functional Brain Imaging on a 3T Phillips scanner. T1-weighted structural images were collected in 150 axial slices with 1 mm isotropic voxels using a magnetization-prepared rapid gradient-echo (MP-RAGE) sequence. Functional BOLD scans were collected in 36 axial slices (2.4 × 2.4 × 3 mm voxels, TR = 2s). Six dummy scans were collected at the beginning of each run but were not saved.

### Analysis

#### Behavioral

We used a mixed-effect logistic regression model to investigate the effect of word category (3 categories: inanimate real word, animate real word, and pseudoword) on in-scanner response accuracy with participant as a random effect. To test statistical significance, we contrasted this model against a null model with the random effect only and no fixed effect. The null model represents the situation in which the performance for all three categories were the same, with only potential idiosyncrasy across participants. Similarly, we used mixed-effect linear regression models to investigate the effect of word category (inanimate real word, animate real word, and pseudoword), uncontracted word length, and word frequency on in-scanner behavioral response times.

#### Whole-cortex univariate analysis

We used FSL, Freesurfer, the HCP workbench, and in-house software to conduct univariate GLM analysis. The cortical surface model for each participant was created using the standard Freesurfer pipeline (Dale, Fischl, & Sereno, 1999; Glasser et al., 2013; Smith et al., 2004). Functional data were motion-corrected, high-pass filtered with a 128s cut-off, and re-sampled to the cortical surface. The surface data were then smoothed with a 6mm FWHM Gaussian kernel and prewhitened to remove temporal autocorrelation. Cerebellar and subcortical structures were excluded.

##### Whole-cortex univariate letter length and word frequency effects

In this univariate GLM analysis, the length of individual inanimate real words in Roman letter spelling (i.e., number of letters in the uncontracted form) and logarithm of word frequency in the COCA database (Davies, 2008) were included as predictors to model their respective effects. Both the log-frequency and the letter length predictors were mean-centered. Four other predictors were included in the model to control for the effect of word categories (inanimate real word, animate real word, and pseudoword) and missed trials in which participants failed to respond.

##### Whole-cortex univariate words vs. pseudoword comparison

In this univariate GLM analysis, for each vertex on the cortical surface, three predictors modeled the effect of inanimate real words, animate real words, and pseudowords, respectively. A separate predictor was added to model missed trials. Since behavioral analysis revealed a significant difference in response time among the three word categories, we included a fifth predictor to model the effect of response time (Yarkoni, Barch, Gray, Conturo, & Braver, 2009).

All predictors modeled only the first three seconds of each trial during which a braille word was presented and were convolved with a canonical hemodynamic response function (Dimsdale-Zucker & Ranganath, 2019). Individual predictors were used to model time points with excessive motion (FDRMS>1.5mm). Within participants, data from different runs were combined using fixed-effects models. Across participants, data were combined using a random-effects model. The resulting maps of p-values underwent cluster-based permutation correction to control the family-wise error rate (FWER). The cluster forming threshold was uncorrected p < 0.01, and the cluster-wise FWER threshold was p < 0.05 corrected.

#### Multi-variate pattern analysis (MVPA)

##### Whole-cortex searchlight SVM decoding: words vs pseudowords

Support vector machine (SVM) decoding implemented in the Python toolbox Scikit-learn was used to distinguish real words from pseudowords based on the spatial activation pattern in each searchlight along the cortex (C.-C. Chang & Lin, 2011; Kriegeskorte, Goebel, & Bandettini, 2006; Pedregosa et al., 2011). A searchlight associated with a vertex on the inflated cortical surface consisted of all the vertices within a circle of 8mm diameter (according to geodesic distance) centered at the vertex (Glasser et al., 2013; Kriegeskorte et al., 2006). Vertices whose corresponding searchlight contained any sub-cortical vertex were excluded.

A GLM was constructed in which each animate, inanimate, and pseudoword was entered as an individual predictor, along with a predictor for missed trials and another predictor for response time. Our first decoding analysis focused on distinguishing real inanimate words from pseudowords, since these are the conditions for which we had the largest amount of data. However, since each inanimate real word appeared 30 times (twice per run), but each pseudoword appeared only twice throughout the experiment, we took several steps to ensure that decoding of inanimate words vs. pseudowords was not driven by different degrees of noise across conditions. First, we matched the number of trials that contributed to the Beta estimate of each word/pseudoword across conditions by extracting one Beta parameter estimate per run for each inanimate real word, for each participant, and extracting one pseudoword Beta for each participant, across runs. This resulted in the same number of trials (two for each entry) contributing to the estimates of Betas for each entry of words and pseudowords. Consequently, in the training dataset for each participant, there was 1 spatial pattern of Beta estimates for each pseudoword and 15 patterns for each inanimate real word (1 pattern from each run). The real word and pseudoword data were separately normalized across items and vertices to ensure that decoding was based on the difference in spatial pattern rather than univariate activation level. Normalization was done by subtracting the mean value and dividing by the standard deviation across patterns and vertices.

Next, we matched the number of training patterns across conditions by splitting the real inanimate word Beta patterns (total of 450) into 15 decoding bins. Each of 15 classifiers was trained to distinguish words and pseudowords using only one of the 15 Beta estimate patterns from each inanimate word, each pattern derived from two trials, and likewise one pattern for each pseudoword, also derived from two trials of that pseudoword. Thus, each classifier had an equivalent amount of data for real and pseudowords. In each of the 15 decoding bins, in each searchlight region, we trained a linear SVM classifier (regularization parameter C = 1) on 90% of the data (27 inanimate words and 27 pseudowords) and tested it on the left out 10% of the data. The decoding accuracy was averaged across 10 train-test splits to derive the accuracy for each decoding bin. Then, the accuracy was averaged across all 15 bins to derive the accuracy for the center vertex of a given searchlight region. Eventually this analysis yielded one whole-brain classification accuracy map for each participant.

We also performed whole-cortex searchlight SVM decoding analysis to distinguish between real animate words and pseudowords. The procedures for this analysis were identical to the inanimate-vs-pseudoword decoding analysis except that decoding attempts were not repeated for 15 decoding bins, because the amount of data contributing to animate words and pseudoword Betas was the same.

We used a permutation and bootstrapping-based method to test the classifier performance against chance (50%) (Elli, Lane, & Bedny, 2019; Schreiber & Krekelberg, 2013; Stelzer, Chen, & Turner, 2013). For each participant, at each vertex, the accuracy value was Fisher-z transformed. We then computed the t-statistics of the group mean (across participants) when tested against chance (which was also Fisher-z transformed). This step resulted in a map of t-statistics. For each participant, we shuffled the word labels (“inanimate word” and “pseudoword”) 200 times, and derived one null accuracy map per shuffle. For each shuffle, we similarly derived across participants a map of t-statistics. Next, in each vertex, we defined the empirical p-value as the probability of observing, in the normal distribution formed by the null values, a t-statistic higher than the actual t-value.

The resulting maps of p-values underwent cluster-based permutation correction to control the family-wise error rate (FWER) (Su, Fonteneau, Marslen-Wilson, & Kriegeskorte, 2012). The cluster forming threshold was uncorrected p < 0.01, and the cluster-wise FWER threshold was p < 0.05 corrected.

### Neural signatures of individual braille words: Split-half MVPA correlation analysis

In a final analysis, we used MVPA to search for neural signatures of individual braille words. Specifically, we conducted a split-half correlation analysis and ROI-based 30-way SVM decoding to investigate whether any cortical areas showed unique spatial patterns of response to specific inanimate real braille words that were distinguishable from all other braille words in our stimulus set. We also performed a similar whole-cortex searchlight MVPA decoding analysis. However, the latter analysis did not yield any significant results and we therefore do not discuss it further here.

In the split-half correlation analysis, we split the data from 15 runs into even runs and the odd runs with run 7 excluded so that both halves contained the same number of runs. We computed one fixed-effect Beta parameter estimation map for each of the 30 words in each half, based on which we created a 30-by-30 similarity matrix between the words in each searchlight. For a diagonal entry (i, i) in such matrix, we computed the Pearson correlation between the spatial pattern of word i in the even half and the pattern of word i in the odd half. For an off-diagonal entry (i, j) in such matrix, we computed the correlation between word i in the even half and word j in the odd half, and the correlation between word j in the even half and word i in the odd half, then took the average of the two correlation values. In each participant, if each word is consistently represented in a searchlight, the spatial patterns of the same word from both halves should be highly correlated, while the patterns of different words should be uncorrelated, leading to the values in the diagonal of the similarity matrix close to 1, and the off-diagonal entries close to 0. We used a bootstrapping method to test this hypothesis. In each searchlight, we computed the mean value across the diagonal entries of the similarity matrix, denoted as M. Then, we computed the mean and standard deviation of the off-diagonal entries in the upper triangle. Next, we computed the z-score of M with respect to the distribution of the off-diagonal values (assuming the off-diagonal values were normally distributed). To test the statistical significance of the resultant z-score in each searchlight, we shuffled all the values in the upper triangle and diagonal entries to create a null similarity matrix and repeated the previous steps to derive a null z-scores. The similarity matrix was shuffled 200 times to derive a null distribution of the z-scores based on which the p-values of the observed z-score was derived. The average z-score map across participants underwent FWER correction with a cluster forming threshold of uncorrected p<0.01 and cluster-wise p<0.05.

## Results

### Behavioral

Participants were highly accurate across conditions, with the highest accuracy for inanimate real words (M=98.8%, SD=2.8%), followed by pseudowords (M=93.8%, SD=4.6%) and finally by animate real words (M=91.1%, SD=11.5%) (mixed-effect logistic regression χ^2^(5)=202.68, p<0.001) (Supplementary Figure 1a).

Participants responded fastest to inanimate real words (M=1.02s, SD=0.38s), followed by animate real words (M=1.55s, SD=0.49s), and slowest for pseudowords (M=1.90s, SD=0.69s) (mixed-effect linear regression χ^2^(2)=3586.1, p<0.001) (Supplementary Figure 1b).

Accuracy for higher frequency words was significantly higher (correlation value M=0.04, SD=0.05; χ^2^(1)=35.16, p<0.001) and response time for higher frequency words was significantly shorter (correlation value M=-0.10, SD=0.04; χ^2^(1)=108.33, p<0.001; mixed-effect linear regression). However, there was no correlation between uncontracted word-length and accuracy (correlation value M=-0.01, SD=0.06; χ^2^(1)=0.07, p=0.79) or response time (correlation value M=-0.02, SD=0.04; χ^2^(1)=1.65, p=0.20). Contracted words whose corresponding uncontracted forms have more letters did not take more time to read than uncontracted words with the same number of cells, nor did they lead to more errors in the lexicality judgement in the current experiment.

### fMRI

#### Neural activity increases in bilateral posterior parietal cortex (PPC) with uncontracted word-length: univariate whole-cortex analysis

In a whole-cortex analysis, we searched for cortical areas where activity increased as the uncontracted forms of words became longer in number of Roman letters, independent of physical word length in braille cells. Bilateral PPC, specifically the supramarginal gyus (SMG) (peak: left -36, -36, 37; right 33, - 44, 40), responded more to words with a larger uncontracted word-length. A small uncontracted word-length responsive cluster was also observed in the right posterior middle frontal gyrus (Figure 1; also see Supplementary Figure 2 for the correlation between uncontracted word length and neural activation in bilateral SMG). No regions increased activity as corresponding uncontracted words became shorter. We conducted two additional whole-cortex univariate analyses where the predictor for uncontracted word-length was replaced by a predictor for number of phonemes and number of syllables, respectively. Despite the high correlation between the numbers of letters, phonemes, and syllables (letters vs phonemes: r = 0.73; letters vs syllables: r = 0.66), we didn’t find an effect of the number of phonemes or syllables under the same threshold and statistical correction as used for number of letters. Nevertheless, due to the high correlation between the numbers of letters and phonemes in English, we cannot unambiguously distinguish between an orthographic effect of underlying uncontracted word length (number of letters) and a phonological effect of the underlying numbers of phonemes or syllables.

**Figure 1:**
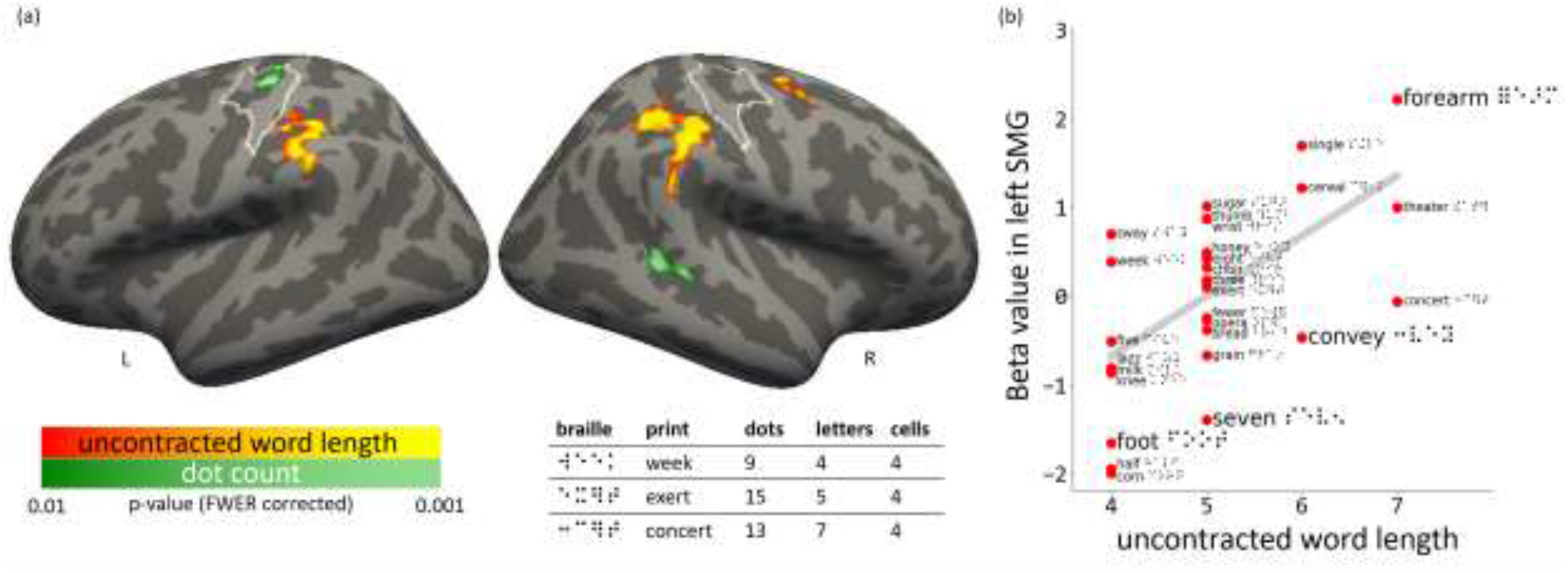
(a) Red/yellow: Regions showing greater response to 4-cell Braille words which contain more letters when transcribed to written English. Dark/bright green: Regions showing greater response to 4-cell Braille words which contain more raised dots. The brain map was cluster-based permutation corrected to control the family-wise error rate (FWER). The cluster forming threshold was uncorrected p < 0.01, and the cluster-wise FWER threshold was p < 0.05. Bottom right: examples of Braille words in this experiment with the least dots (week) and the most dots (exert); “week” is also one of the shortest words in Roman spelling (4 letters), while “concert” is one of the longest (7 letters). White outlines mark the hand S1/M1 region. (b) The correlation between word length in uncontracted Roman spelling and neural activation in the left supramarginal gyri (SMG), measured with the beta estimation for the activation level of each word. For illustration purpose, in the left SMG, for each word, we averaged the beta value across the vertices and across participants. Then, we correlated the average beta values with the word lengths.

For the words used in this experiment, another variable that was moderately, but significantly, correlated with uncontracted word-length was the number of raised dots in a word, because the cells corresponding to contractions are, on average, more likely to have a larger number of dots than those corresponding to individual letters. For the 30 inanimate real words used in this experiment, the correlation between uncontracted word length and dot count is r=0.44. We conducted an additional whole-cortex univariate analyses in which the predictor for uncontracted word-length was replaced by a predictor for dot count. We observed an increase in activity as a function of dot count in the hand region of the left primary somatosensory cortex (S1). Figure 1 shows this activation relative to an outline of the hand region for left sensorimotor cortex from Neurosynth (Loiotile, Lane, Omaki, & Bedny, 2020; Tian et al., 2022). This effect was clearly superior and anterior and disjoint from the cluster showing the uncontracted word-length effect. A dot-number effect was also observed in the right superior temporal sulcus.

#### Neural effects of word frequency are differently localized from effects of uncontracted word-length: univariate whole-cortex analysis

Responses that showed sensitivity to word frequency were localized to different cortical areas from those that exhibited sensitivity to of uncontracted word-length (above). We observed larger responses to low frequency words in the left IFS, left superior frontal gyrus (SFG), right fusiform gyrus, and right anterior inferior/middle temporal gyri (Supplementary Figure 3). No regions responded more to higher compared to lower frequency words. No word frequency effects were observed in the SMG or anywhere in the PPC.

#### Univariate differences between words vs pseudowords in whole-cortex analysis

We observed larger responses to pseudowords than words (inanimate) bilaterally in PPC (intraparietal sulcus (IPS) and supramarginal gyrus (SMG)) and in inferior frontal cortex (inferior frontal sulcus (IFS)/precentral sulcus (PCS), anterior insula) (Figure 2a). Larger responses to pseudowords than words were also observed in ventral occipto-temporal cortex vOTC (fusiform gyrus) bilaterally. vOTC responses were located medial and anterior to the location of the VWFA region as defined based on a meta-analysis carried out by Jobard et al. (2003). Some responses were also observed lateral to this VWFA location, extending into the inferior temporal gyrus and the middle occipital sulcus on the lateral surface. Larger responses to pseudowords were also observed in dorsal occipital and parieto-occipital areas as well as in right primary visual cortex. We observed a qualitatively similar but weaker effects when comparing pseudowords to animate real words (for a complete list of regions in both contrasts see Table 5).

**Figure 2:**
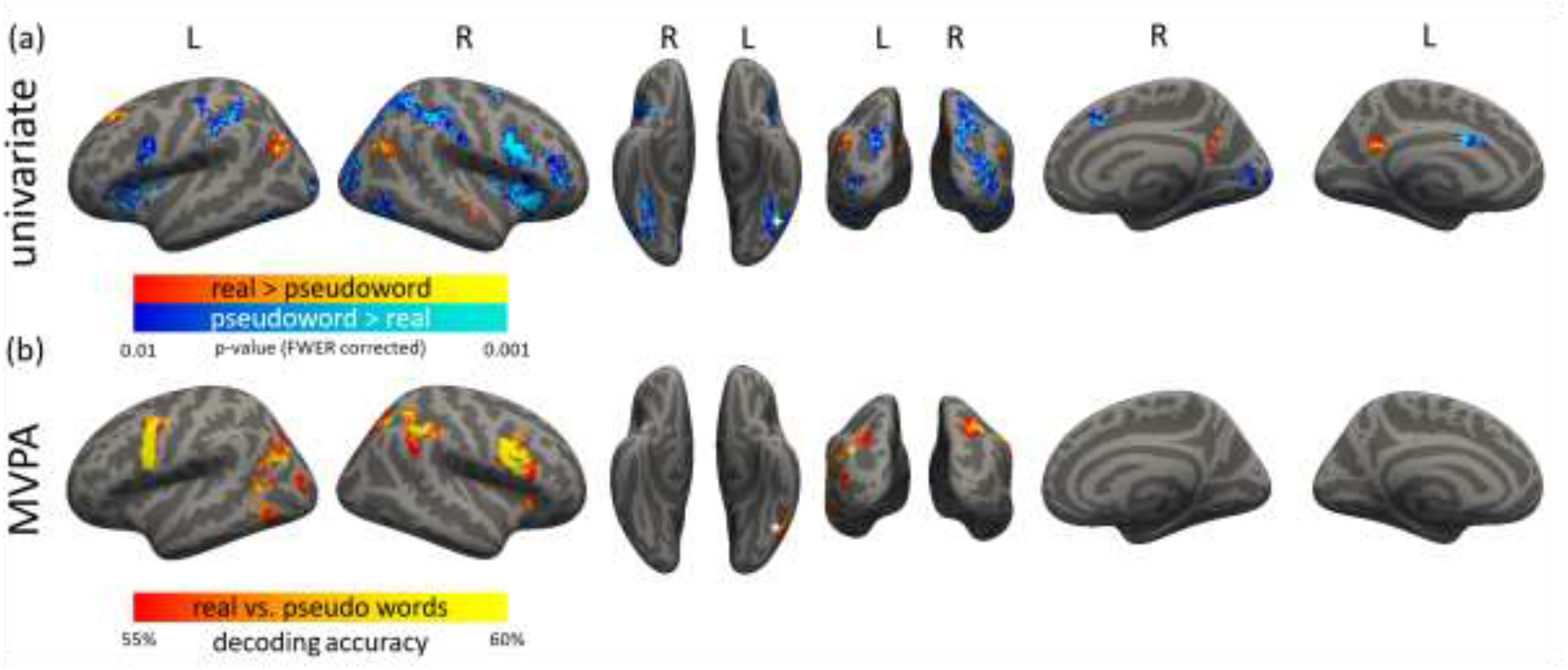
Differences in the neural response to real words and pseudowords. (a) univariate contrasts. Warm color: inanimate real words > pseudowords. Cool color: pseudowords > inanimate real words. (b) Inanimate real word vs pseudoword MVPA decoding accuracy. Chance level of decoding accuracy = 50%. Both maps underwent cluster-based permutation correction to control the family-wise error rate (FWER). The cluster forming threshold was uncorrected p < 0.01, and the cluster-wise FWER threshold was p < 0.05. The peak of the VWFA reported by Jobard et al. (2003) is marked with the white cross.

**Table 5.**
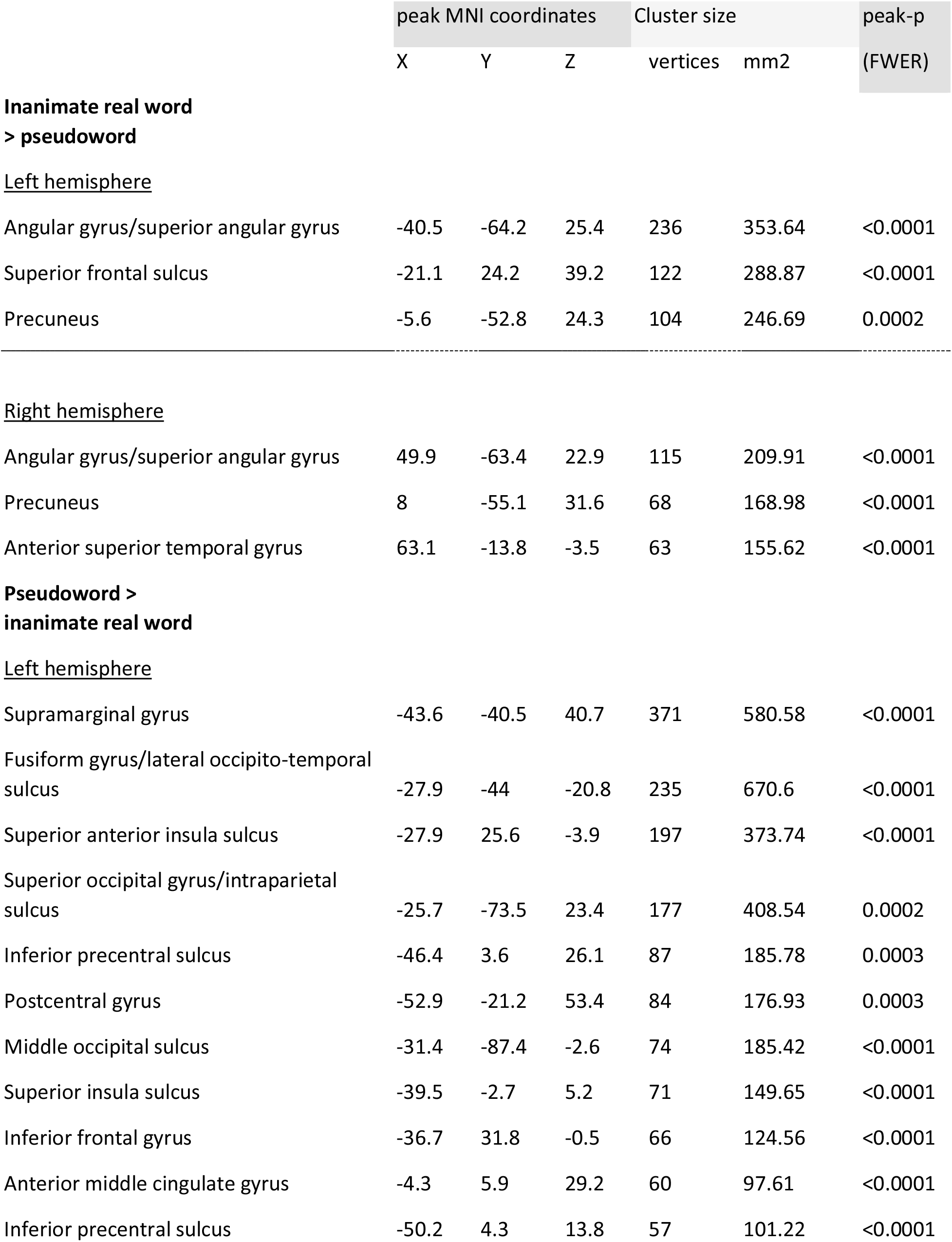
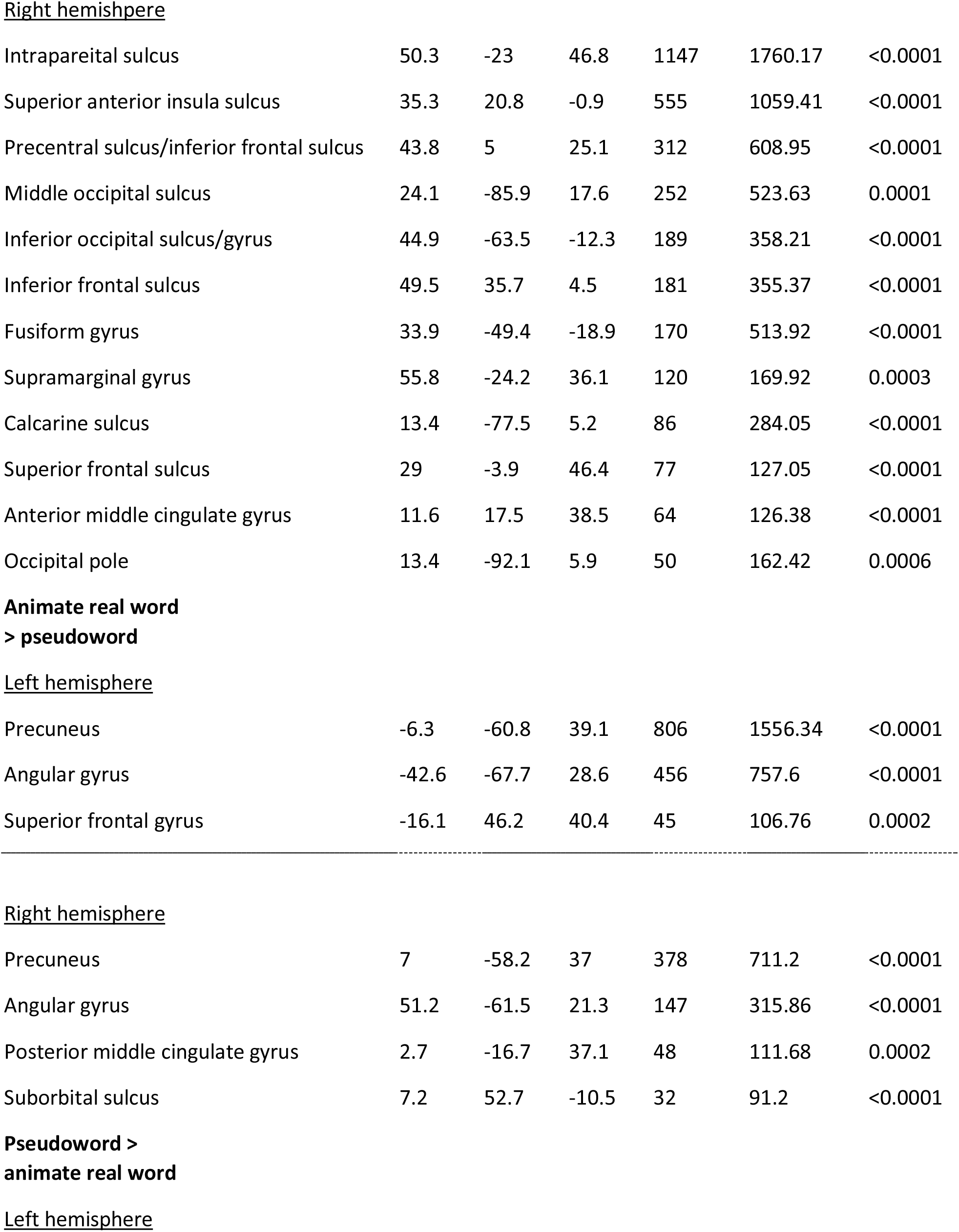
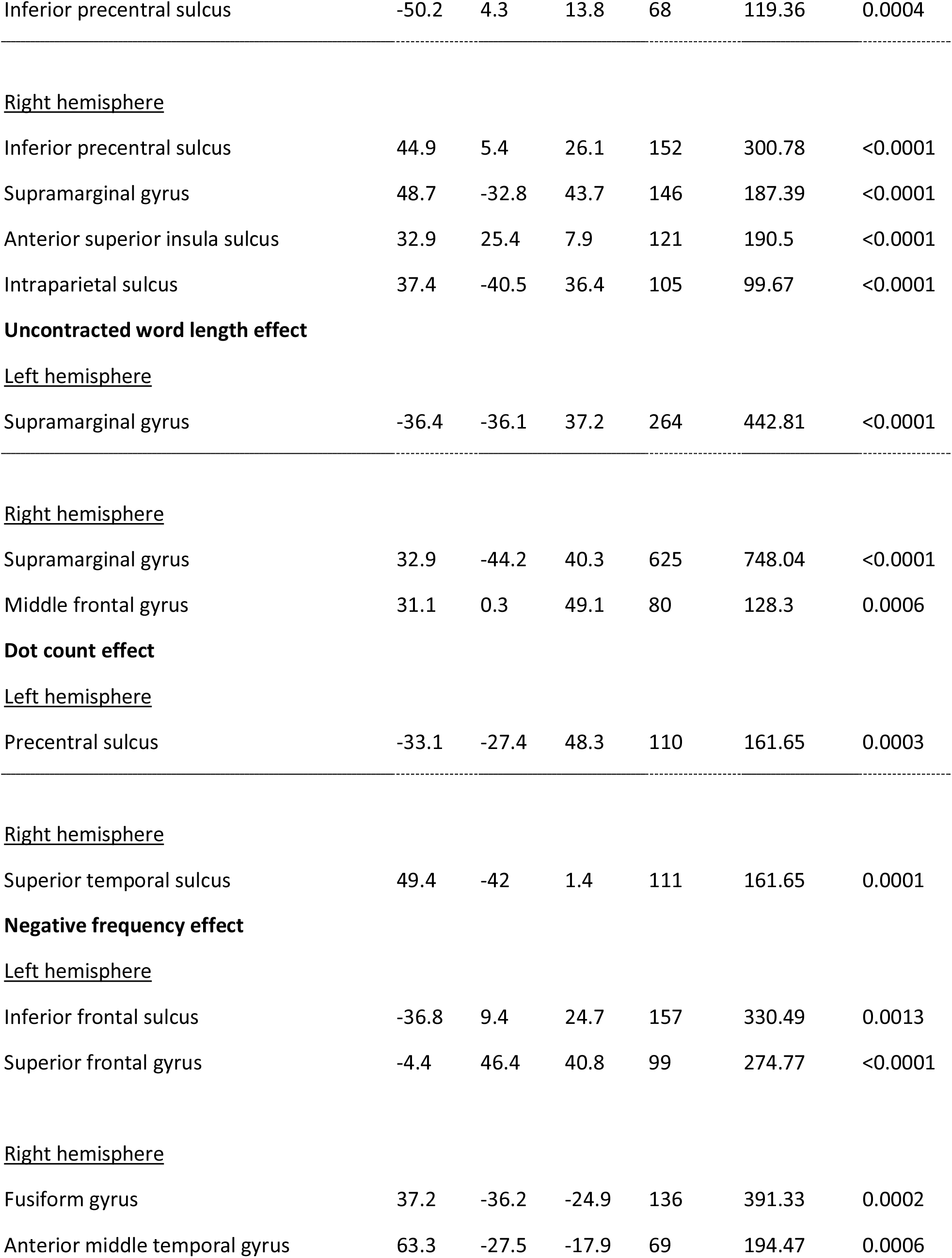
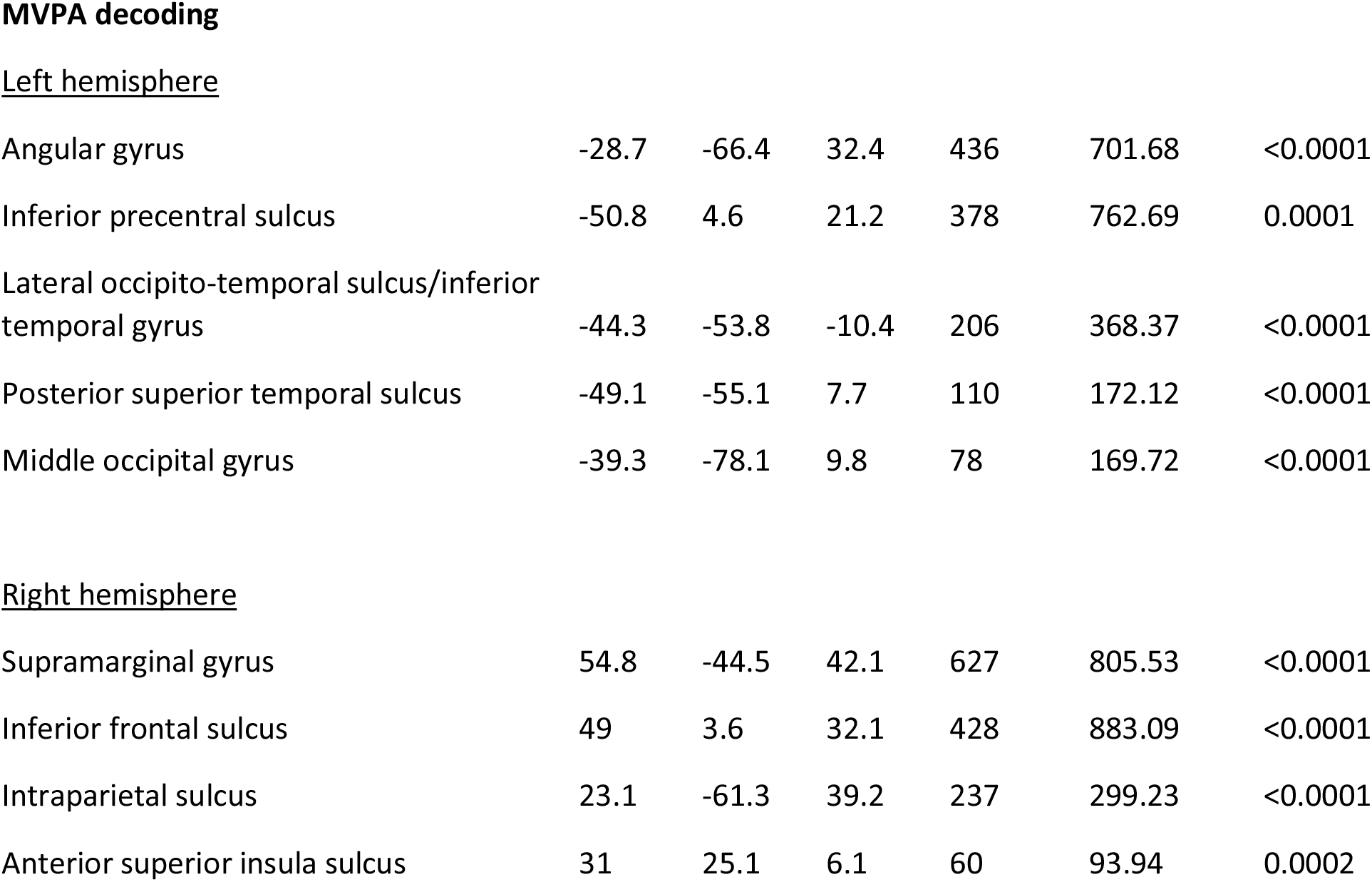
List of activated clusters

|  | peak MNI coordinates |  |  | Cluster size |  | peak-p |
| --- | --- | --- | --- | --- | --- | --- |
|  | X | Y | Z | vertices | mm2 | (FWER) |
| **Inanimate real word** |  |  |  |  |  |  |
| **> pseudoword** |  |  |  |  |  |  |
| <u>Left hemisphere</u> |  |  |  |  |  |  |
| Angular gyrus/superior angular gyrus | -40.5 | -64.2 | 25.4 | 236 | 353.64 | <0.0001 |
| Superior frontal sulcus | -21.1 | 24.2 | 39.2 | 122 | 288.87 | <0.0001 |
| Precuneus | -5.6 | -52.8 | 24.3 | 104 | 246.69 | 0.0002 |
| <u>Right hemisphere</u> |  |  |  |  |  |  |
| Angular gyrus/superior angular gyrus | 49.9 | -63.4 | 22.9 | 115 | 209.91 | <0.0001 |
| Precuneus | 8 | -55.1 | 31.6 | 68 | 168.98 | <0.0001 |
| Anterior superior temporal gyrus | 63.1 | -13.8 | -3.5 | 63 | 155.62 | <0.0001 |
| **Pseudoword >** |  |  |  |  |  |  |
| **inanimate real word** |  |  |  |  |  |  |
| <u>Left hemisphere</u> |  |  |  |  |  |  |
| Supramarginal gyrus | -43.6 | -40.5 | 40.7 | 371 | 580.58 | <0.0001 |
| Fusiform gyrus/lateral occipito-temporal sulcus | -27.9 | -44 | -20.8 | 235 | 670.6 | <0.0001 |
| Superior anterior insula sulcus | -27.9 | 25.6 | -3.9 | 197 | 373.74 | <0.0001 |
| Superior occipital gyrus/intraparietal sulcus | -25.7 | -73.5 | 23.4 | 177 | 408.54 | 0.0002 |
| Inferior precentral sulcus | -46.4 | 3.6 | 26.1 | 87 | 185.78 | 0.0003 |
| Postcentral gyrus | -52.9 | -21.2 | 53.4 | 84 | 176.93 | 0.0003 |
| Middle occipital sulcus | -31.4 | -87.4 | -2.6 | 74 | 185.42 | <0.0001 |
| Superior insula sulcus | -39.5 | -2.7 | 5.2 | 71 | 149.65 | <0.0001 |
| Inferior frontal gyrus | -36.7 | 31.8 | -0.5 | 66 | 124.56 | <0.0001 |
| Anterior middle cingulate gyrus | -4.3 | 5.9 | 29.2 | 60 | 97.61 | <0.0001 |
| Inferior precentral sulcus | -50.2 | 4.3 | 13.8 | 57 | 101.22 | <0.0001 |
| Right hemisphere |  |  |  |  |  |  |
| Intrapareital sulcus | 50.3 | -23 | 46.8 | 1147 | 1760.17 | <0.0001 |
| Superior anterior insula sulcus | 35.3 | 20.8 | -0.9 | 555 | 1059.41 | <0.0001 |
| Precentral sulcus/inferior frontal sulcus | 43.8 | 5 | 25.1 | 312 | 608.95 | <0.0001 |
| Middle occipital sulcus | 24.1 | -85.9 | 17.6 | 252 | 523.63 | 0.0001 |
| Inferior occipital sulcus/gyrus | 44.9 | -63.5 | -12.3 | 189 | 358.21 | <0.0001 |
| Inferior frontal sulcus | 49.5 | 35.7 | 4.5 | 181 | 355.37 | <0.0001 |
| Fusiform gyrus | 33.9 | -49.4 | -18.9 | 170 | 513.92 | <0.0001 |
| Supramarginal gyrus | 55.8 | -24.2 | 36.1 | 120 | 169.92 | 0.0003 |
| Calcarine sulcus | 13.4 | -77.5 | 5.2 | 86 | 284.05 | <0.0001 |
| Superior frontal sulcus | 29 | -3.9 | 46.4 | 77 | 127.05 | <0.0001 |
| Anterior middle cingulate gyrus | 11.6 | 17.5 | 38.5 | 64 | 126.38 | <0.0001 |
| Occipital pole | 13.4 | -92.1 | 5.9 | 50 | 162.42 | 0.0006 |
| Precuneus | -6.3 | -60.8 | 39.1 | 806 | 1556.34 | <0.0001 |
| Angular gyrus | -42.6 | -67.7 | 28.6 | 456 | 757.6 | <0.0001 |
| Superior frontal gyrus | -16.1 | 46.2 | 40.4 | 45 | 106.76 | 0.0002 |
| Right hemisphere |  |  |  |  |  |  |
| Precuneus | 7 | -58.2 | 37 | 378 | 711.2 | <0.0001 |
| Angular gyrus | 51.2 | -61.5 | 21.3 | 147 | 315.86 | <0.0001 |
| Posterior middle cingulate gyrus | 2.7 | -16.7 | 37.1 | 48 | 111.68 | 0.0002 |
| Suborbital sulcus | 7.2 | 52.7 | -10.5 | 32 | 91.2 | <0.0001 |
| Inferior precentral sulcus | -50.2 | 4.3 | 13.8 | 68 | 119.36 | 0.0004 |
| <u>Right hemisphere</u> |  |  |  |  |  |  |
| Inferior precentral sulcus | 44.9 | 5.4 | 26.1 | 152 | 300.78 | <0.0001 |
| Supramarginal gyrus | 48.7 | -32.8 | 43.7 | 146 | 187.39 | <0.0001 |
| Anterior superior insula sulcus | 32.9 | 25.4 | 7.9 | 121 | 190.5 | <0.0001 |
| Intraparietal sulcus | 37.4 | -40.5 | 36.4 | 105 | 99.67 | <0.0001 |
| **Uncontracted word length effect** |  |  |  |  |  |  |
| <u>Left hemisphere</u> |  |  |  |  |  |  |
| Supramarginal gyrus | -36.4 | -36.1 | 37.2 | 264 | 442.81 | <0.0001 |
| <u>Right hemisphere</u> |  |  |  |  |  |  |
| Supramarginal gyrus | 32.9 | -44.2 | 40.3 | 625 | 748.04 | <0.0001 |
| Middle frontal gyrus | 31.1 | 0.3 | 49.1 | 80 | 128.3 | 0.0006 |
| **Dot count effect** |  |  |  |  |  |  |
| <u>Left hemisphere</u> |  |  |  |  |  |  |
| Precentral sulcus | -33.1 | -27.4 | 48.3 | 110 | 161.65 | 0.0003 |
| <u>Right hemisphere</u> |  |  |  |  |  |  |
| Superior temporal sulcus | 49.4 | -42 | 1.4 | 111 | 161.65 | 0.0001 |
| **Negative frequency effect** |  |  |  |  |  |  |
| <u>Left hemisphere</u> |  |  |  |  |  |  |
| Inferior frontal sulcus | -36.8 | 9.4 | 24.7 | 157 | 330.49 | 0.0013 |
| Superior frontal gyrus | -4.4 | 46.4 | 40.8 | 99 | 274.77 | <0.0001 |
| <u>Right hemisphere</u> |  |  |  |  |  |  |
| Fusiform gyrus | 37.2 | -36.2 | -24.9 | 136 | 391.33 | 0.0002 |
| Anterior middle temporal gyrus | 63.3 | -27.5 | -17.9 | 69 | 194.47 | 0.0006 |
| Left hemisphere |  |  |  |  |  |  |
| Angular gyrus | -28.7 | -66.4 | 32.4 | 436 | 701.68 | <0.0001 |
| Inferior precentral sulcus | -50.8 | 4.6 | 21.2 | 378 | 762.69 | 0.0001 |
| Lateral occipito-temporal sulcus/inferior temporal gyrus | -44.3 | -53.8 | -10.4 | 206 | 368.37 | <0.0001 |
| Posterior superior temporal sulcus | -49.1 | -55.1 | 7.7 | 110 | 172.12 | <0.0001 |
| Middle occipital gyrus | -39.3 | -78.1 | 9.8 | 78 | 169.72 | <0.0001 |
| Right hemisphere |  |  |  |  |  |  |
| Supramarginal gyrus | 54.8 | -44.5 | 42.1 | 627 | 805.53 | <0.0001 |
| Inferior frontal sulcus | 49 | 3.6 | 32.1 | 428 | 883.09 | <0.0001 |
| Intraparietal sulcus | 23.1 | -61.3 | 39.2 | 237 | 299.23 | <0.0001 |
| Anterior superior insula sulcus | 31 | 25.1 | 6.1 | 60 | 93.94 | 0.0002 |

Interestingly, the parietal and frontal regions showing responses to uncontracted word-length overlapped with parietal and frontal regions more active for pseudowords relative to real words (Figure 2; see Figure 3 for the overlap), despite the fact that pseudowords were not included in the uncontracted word-length analysis. This provides further evidence for the possibility that these parietal regions play a role in form-based orthographic braille processing, perhaps the conversion from contracted to uncontracted Roman spellings and/or from graphemes to phonemes.

**Figure 3:**
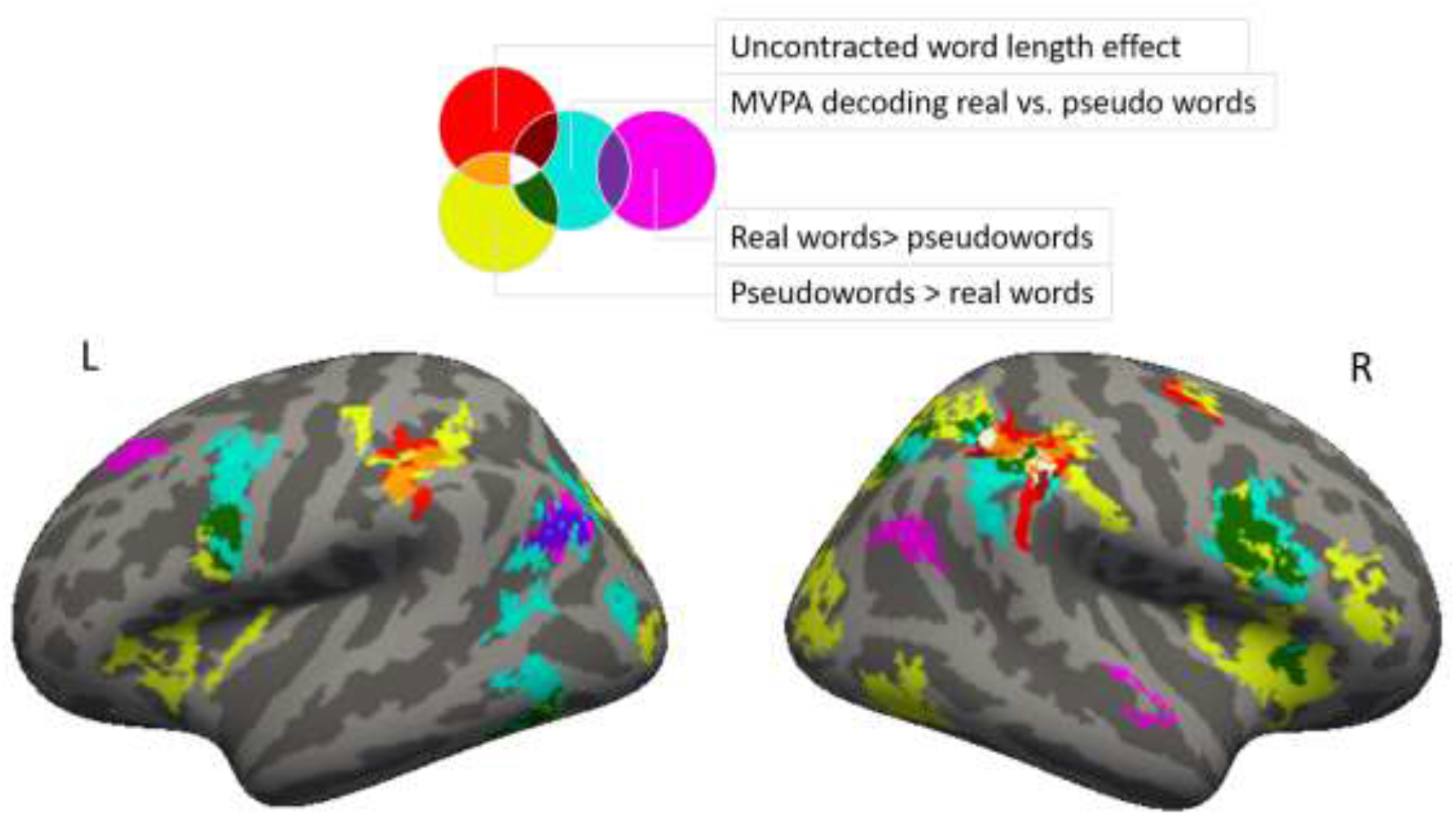
Overlap between four maps: univariate inanimate real words > pseudowords (R>P, magenta), univariate pseudowords > inanimate real words (P>R, yellow), MVPA decoding (MVPA, cyan), uncontracted word length effect (LEN, red). Purple: overlap between R>P and MVPA. Green: overlap between P>R and MVPA. Orange: overlap between P>R and LEN. Dark red: overlap between MVPA and LEN. White: overlap between P>R, MVPA, and LEN.

The univariate contrast of words (inanimate) vs. pseudowords revealed greater activation for words in areas associated with semantic processing, including medially, the bilateral precuneus (PC) and on the lateral surface bilateral temporo-parietal cortex (angular gyri, AG) (Figure 2a). Small clusters were also observed in the right anterior superior temporal gyrus (STG) and left superior frontal sulcus (SFS).

#### Words vs pseudowords MVPA whole-cortex analysis

Words and pseudowords produced different MVPA patterns in right PPC (SMG, extending into the AG and the IPS) and inferior frontal cortex (in bilateral IFS/PCS and right anterior insula) (Figure 2b). These areas overlapped with univariate pseudoword responses as well as to some extent with uncontracted word-length responsive areas. There was also some overlap with responses to words in the left temporo-parietal junction (AG, extending ventrally into the posterior superior temporal sulcus (STS) dorsally into the posterior IPS). Additional areas of significant decoding that did not overlap with the univariate analysis were observed in left occipital and occipito-temporal cortices (posterior inferior temporal sulcus and middle occipital sulcus/gyrus).

#### Exploratory analyses of univariate and multivariate effects in the vOTC at a lower threshold

Given the involvement of VWFA in the processing of orthography in the sighted and prior reports of responses to braille in this region, we searched for effects in the vOTC at an exploratory threshold to see if effects in the classic VWFA location would emerge (p-values less than 0.05, uncorrected). At this exploratory threshold, in the pseudowords > inanimate real words univariate contrast, activation differences were observed throughout the left vOTC, including but not limited to the VWFA location. In the map of MVPA decoding of pseudowords vs. words and the map of uncontracted word-length effect, even at this exploratory threshold, the involvement of the left vOTC was restricted to the inferior temporal sulcus and anterior fusiform gyrus. In the map of the negative frequency effect (higher activation for lower frequency words), the lateral and medial portion of the left vOTC were identified, but in both maps, the peak of the VWFA as reported in the meta-analysis by Jobard et al.(2003) was not recruited (Supplementary Figure 4).

#### Neural signatures of individual braille words: Split-half MVPA correlation analysis

We attempted to identify cortical areas containing information about individual word identities, relative to all other words in our stimulus set. In a whole-cortex searchlight split-half analysis that searched for cortical regions where the similarity between a word and itself was greater than between that word and all other words in the stimulus set, we observed small significant clusters in bilateral inferior precentral sulcus, bilateral posterior intraparietal sulcus, bilateral precuneus, and left posterior superior temporal sulcus. Except for the clusters in bilateral precuneus, these clusters overlapped with the regions that distinguished between inanimate words and pseudowords in the whole-cortex searchlight SVM decoding analysis (Supplementary Figure 5).

## Discussion

We observed orthographic reading-related effects for braille that were separable from low-level tactile and high-level semantic processes in the posterior parietal cortices (PPC). Activation in bilateral SMG (left -36, -36, 37; right 33, -44, 40) was positively correlated with the lengths of uncontracted words corresponding to the contracted braille words that the subjects were actually reading. Braille words consisting of the same number of cells but with more letters in their corresponding uncontracted forms (e.g., “milk”, ⠍⠊⠇⠅ vs. “concert” ⠒⠉⠻⠞) produced higher SMG activity. Neither the number of syllables, phonemes, or dots predicted this SMG effect. By contrast, the hand region of primary somatosensory cortex was sensitive to total dot number per word, a proxy for the amount of somatosensory stimulation. In the current study and in prior work, contractions did not slow down proficient readers (Millar, 1997). Thus, rather than reflecting reading difficulty, the effect of the uncontracted word length is likely to reflect the retrieval of the sublexical units in braille (Fischer-Baum & Englebretson, 2016). One possibility is that in addition to understanding contractions as stand-alone symbols, letters represented by the contractions are retrieved and reorganized according to the morphological structure of the word (Fischer-Baum & Englebretson, 2016).

In addition to showing an uncontracted word length effect, bilateral SMG also showed other signatures of orthographic processing. The SMG showed greater activation for pseudowords than real words and right SMG showed above-chance MVPA decoding of real and pseudowords, even though the pseudowords did not differ from real words in low-level tactile properties. In the whole-cortex analyses, only the right SMG exhibited all three effects: the uncontracted word-length effect, the pseudoword preference, and differentiation of real versus pseudowords in MVPA. The dorsal occipito-parietal, and bilateral prefrontal cortices also differentiated pseudowords from real words but did not show an uncontracted word-length effect, these regions may also have orthographic functions. Together these findings suggest that PPC, and the SMG in particular, plays a role in orthographic processing during braille reading.

An intriguing possibility is that the PPC becomes recruited for braille orthography because of its role in high-level tactile texture and shape perception, analogous to the vOTC’s role in visual object recognition (Burton, MacLeod, Videen, & Raichle, 1997; Chivukula et al., 2021; Debowska et al., 2016; Ro, Wallace, Hagedorn, Farne, & Pienkos, 2004; Sathian, 2016; Stilla, Deshpande, LaConte, Hu, & Sathian, 2007). The SMG is involved in tactile pattern perception and active touch (Bodegård, Geyer, Grefkes, Zilles, & Roland, 2001; Li Hegner, Lee, Grodd, & Braun, 2010). Damage to the PPC can cause tactile agnosia: the inability to recognize or name shapes from touch, the in the absence of low-level somatosensory deficits (Bohlhalter, Fretz, & Weder, 2002; Veronelli, Ginex, Dinacci, Cappa, & Corbo, 2014). After sighted individuals received braille training for three-weeks, structural and functional changes are found in the PPC, in addition to the primary somatosensory cortex (Debowska et al., 2016). The PPC is also structurally connected to somatosensory and language network (Burks et al., 2017; Catani, Jones, & Ffytche, 2005; Frey, Campbell, Pike, & Petrides, 2008; Margulies & Petrides, 2013; Mohan, de Haan, Mansvelder, & de Kock, 2018; Parker et al., 2005; Save & Poucet, 2009). According to this hypothesis, the PPC may play a role in braille word recognition analogous to that of the visual word form area (VWFA) in visual print reading.

Another possibility is that the PPC contributes to braille reading because of its role in orthographic working memory or grapheme to phoneme conversion, analogous to its role in sighted readers (Vogel et al., 2013; Vogel et al., 2012; Vogel, Petersen, & Schlaggar, 2014) and spellers (Graves et al., 2010; Jobard et al., 2003; Purcell, Rapp, & Martin, 2021; Purcell et al., 2011; Rapp & Dufor, 2011; Stoeckel, Gough, Watkins, & Devlin, 2009).

In addition to effects in parietal and prefrontal cortices, we also observed univariate and MVPA sensitivity to pseudowords in the vOTC and the lateral inferior temporal gyrus, consistent with the idea that parts of the ventral stream play a role in braille processing. However, unlike previous studies that contrasted braille against lower-level control conditions in blind readers, the vOTC effects we observed did not show a clear focal peak at the canonical anatomical location associated with the VWFA (−42, -57, -15) (Chen et al., 2019; Cohen et al., 2002; Kronbichler et al., 2004; Vogel et al., 2012). The VWFA may in fact be the peak of linguistic rather than orthographic responses during braille reading, while more medial parts of vOTC specifically respond to orthography. Medial parts of the vOTC are connected with parietal circuits, including SMG, as well as dorsal occipital cortex (Bouhali et al., 2019; Jitsuishi et al., 2020; Leo et al., 2012; Moulton et al., 2019). This connectivity could convey orthographic information to vOTC.

Notably, most of the orthographic responses observed in the current study were bilateral and, if anything, somewhat stronger in the right hemisphere. This is different from the left-lateralized responses to orthography generally observed during visual print reading by sighted people and may be related to changes in spoken language lateralization in the blind (Lane et al., 2016; Röder, Stock, Bien, Neville, & Rösler, 2002).

Finally, we were able to decode individual braille words based on patterns of activity in prefrontal, parieto-occipital and occipito-temporal areas. Notably these effects fell outside of S1, suggesting decoding based on something other than low-level sensory properties. This is unsurprising since previous studies of S1 have only decoded responses to tactile stimulation of different positions along the length of the finger at higher field strengths, and the differences amongst braille words are far subtler (Sanchez-Panchuelo et al., 2012). Even outside of S1, effects were weak and only partially overlapping with the other MVPA and univariate effects. These decoding results serve as a proof of principle that braille words can be decoded based on neural activity patterns, despite their high sensory similarity to each other and the dynamic nature of touch, but also suggest that these signatures are difficult to detect with conventional MVPA analyses.

In summary, we identified several neural signatures of orthographic processing in braille reading. Posterior-parietal and parieto-occipital cortices, including specifically the SMG, are sensitive to form-based orthographic properties of braille. The SMG showed sensitivity to uncontracted word length when participants read contracted braille, a possible signature of processing a dual orthographic code.

Behavioral data, experiment stimuli, and the script for the generation and presentation of the stimuli are available in an OSF repository: https://osf.io/tnbd5/

## Acknowledgements

We are grateful to the participants and the Baltimore blind community without whose support this research would not be possible. We thank Dr. Judy Kim and Lindsay Yazzolino for input on the design of this project; Brianna Alheimer for help with data collection; Dr. Robert Englebretson for valuable intellectual input regarding earlier drafts of this article

