## Supplemental figures for "Neural signatures of reading-related orthographic processing in braille"

PNAS

### Supporting Information for
Neural signatures of reading-related orthographic processing in English braille

Yun-Fei Liu, Brenda Rapp, Marina Bedny

Yun-Fei Liu


**This PDF file includes:**

Figures S1 to S3

**Other supporting materials for this manuscript include the following:**

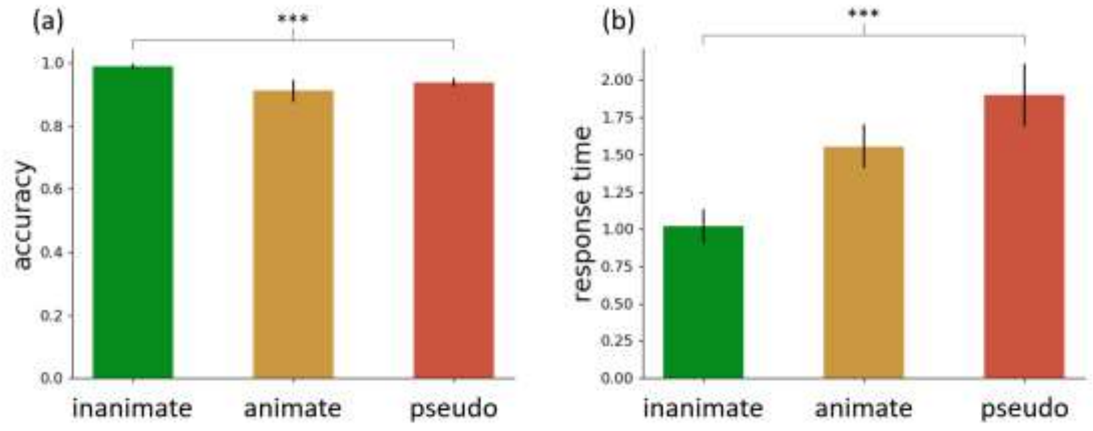


**Fig. S1.** Behavioral results of the fMRI task. (a) Accuracy. (b) Response time. ***p<0.001

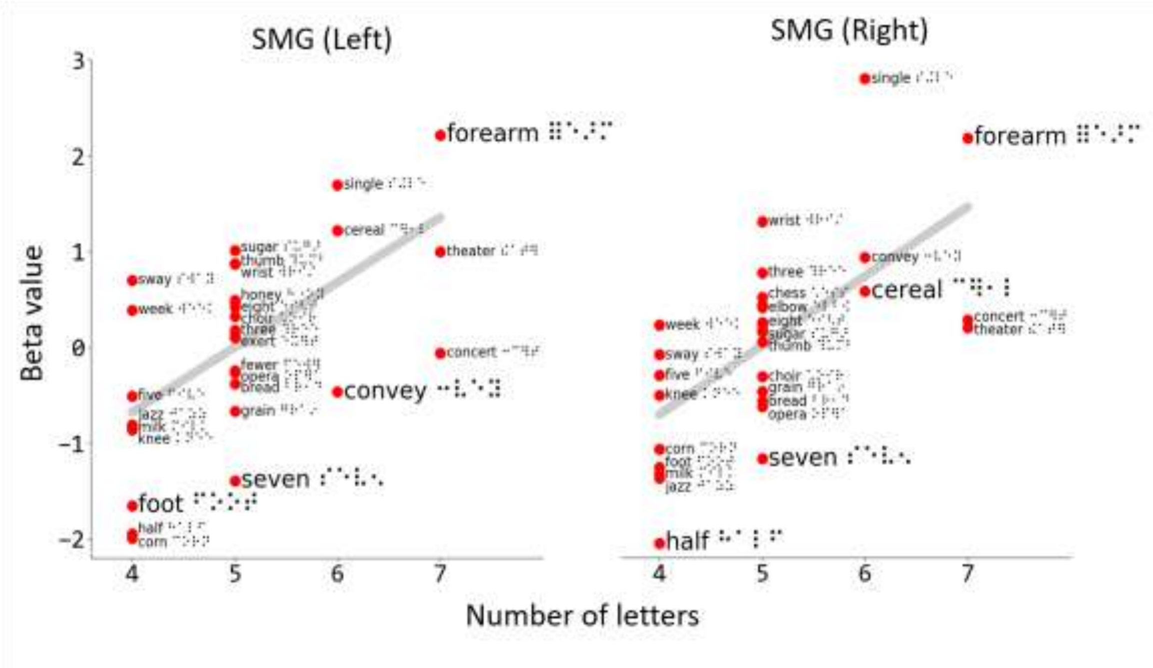


**Fig. S2.** The correlation between word length in uncontracted Roman spelling and neural activation in bilateral supramarginal gyri (SMG). For illustration purpose, in the SMG in either hemisphere, for each word, we averaged the beta value across the vertices and across participants. Then, we correlated the average beta values with the word lengths.

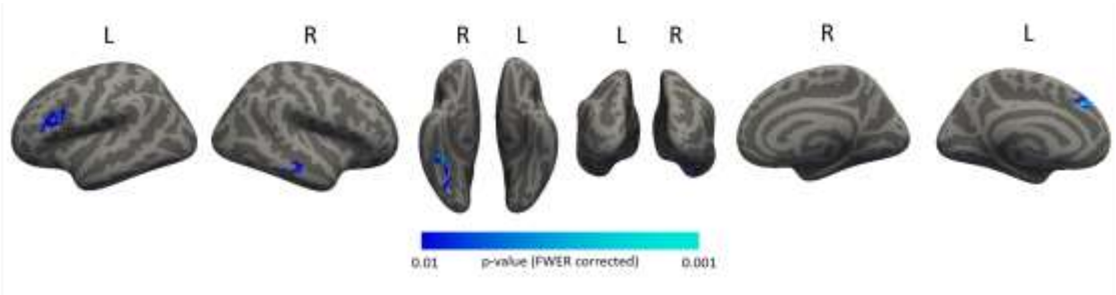


**Fig. S3.** The negative effect of log word frequency, with less activity for higher frequency words. The map was cluster-based permutation corrected to control the family-wise error rate (FWER). The cluster forming threshold was uncorrected $p < 0.01$, and the cluster-wise FWER threshold was $p < 0.05$

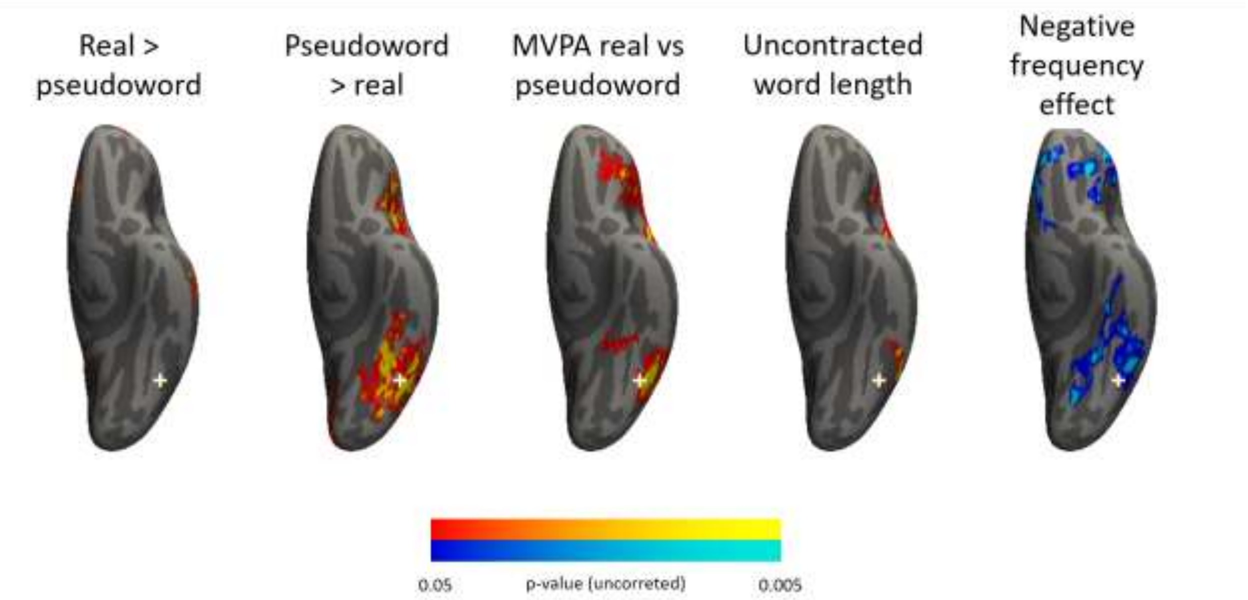


**Fig. S4:** Ventral view of the left hemisphere in five maps in which exploratory thresholds (uncorrected p<0.05) were applied. From left to right: inanimate real word > pseudoword contrast, pseudoword > inanimate real word contrast, inanimate real word vs pseudoword MVPA decoding, letter length effect, negative frequency effect. Only clusters with greater than 50 vertices are shown. No permutation-based cluster correction was performed.

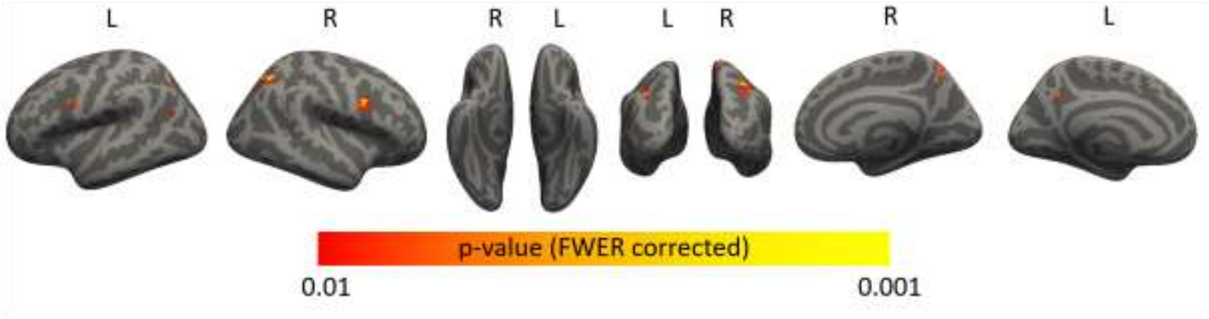


**Fig. S5:** Split-half MVPA searchlight correlation analysis. The brain map shows cortical regions where , averaged across all inanimate real words, the similarity between any inanimate real word and itself was greater than between that word and all other inanimate real words in the stimulus set . The map underwent cluster-based permutation correction to control the family-wise error rate (FWER). The cluster forming threshold was uncorrected $p < 0.01$, and the cluster-wise FWER threshold was $p < 0.05$.
